# Secondary Structure Diversity of the Mitochondrial Small-Subunit rRNA in Porifera

**DOI:** 10.64898/2026.08.28.747467

**Authors:** Zhou Yunqian, Gong Lin, Niu Gengyun, Shi Haihe, Li Xinzheng, Robin Gutell, Wei Meicai

## Abstract

Animal mitochondrial rRNAs are commonly viewed as structurally reduced, yet sponge mt SSU rRNAs range from compact to highly expanded structures. Using nine conserved structural anchors, we compared 216 taxonomically resolved records from four classes and 22 orders, including 16 freshwater Spongillida and 200 marine sponges. Twelve homologous hypervariable substructures were coded as structural types, and their ordered combinations as composite types. We identified 38 structural types and 62 composite types across molecules ranging from 853 to 2,019 nt. Hexactinellida and freshwater Spongillida were each uniform for a distinct composite type but differed markedly in overall structure: hexactinellid mt SSU rRNAs were compact, whereas those of Spongillida were long and contained four to five candidate insertion regions. These results show that a conserved scaffold can accommodate extensive lineage-associated structural variation and provide a practical framework for comparing highly divergent mitochondrial rRNAs.

## 1. Introduction

Mitochondrial rRNAs are commonly described as undergoing reduction during evolution. In animals, the loss of rRNA mass is often accompanied by an increased contribution of mitoribosomal proteins, forming the classical framework of rRNA reduction and protein compensation (van der Sluis et al., 2015). However, this broad tendency does not encompass the full structural diversity of animal mt SSU rRNAs, which range from exceptionally short to highly expanded forms. For example, the demosponge *Suberites domuncula* has a 1,833-nt *rns* gene (Lukić-Bilela et al., 2008), whereas the ctenophore *Mnemiopsis leidyi* retains a highly reduced molecule of only 368 nt (Pett et al., 2011). Thus, the key question is no longer simply whether mitochondrial rRNA is reduced, but which structural elements remain comparable and where lineage-associated remodeling occurs.

A recent large-scale comparison of vertebrate mt SSU rRNAs revealed a broadly conserved canonical structure together with rarer, lineage-restricted types defined mainly by the loss of particular stems (Li et al., 2025). Sponges present a different and complementary problem. Their mt SSU rRNAs include both compact forms and pronounced expansions, with some sequences exceeding the length of the approximately 1.5-kb bacterial SSU rRNA of *Escherichia coli*. Sponge mitochondrial genomes also vary substantially in organization, genetic code, RNA processing, and repetitive-element content (Lavrov et al., 2005, 2013; Wang & Lavrov, 2007). Porifera therefore provides an informative system for examining how a broadly conserved rRNA scaffold accommodates not only reduction but also extensive lineage-associated expansion.

Comparative analysis of rRNA rests on two complementary foundations: rRNA sequences retain phylogenetic information across deep evolutionary distances (Woese, 1987), whereas covariation among homologous positions provides evidence for conserved base pairing and higher-order structure (Gutell et al., 1994). Secondary structures can therefore be inferred by aligning homologous sequences, identifying conserved base pairs, and evaluating proposed stems through compensatory substitutions (Cannone et al., 2002; Gutell et al., 2002). In sponge mt SSU rRNAs, however, long insertions and low sequence identity make primary-sequence alignment alone insufficient for establishing positional homology across classes. Conserved structural anchors do not by themselves demonstrate homology throughout the molecule; rather, repeatedly identifiable stems and loops provide stable boundaries for the adjacent variable regions. Defining these boundaries reduces positional ambiguity and allows the intervening substructures to be compared within a common structural framework. This anchor-first strategy is therefore particularly important for cross-class comparisons of highly divergent sponge sequences.

Building on the comparative framework previously applied to vertebrates (Li et al., 2025), but addressing the contrasting prominence of structural expansion in sponges, here we establish a set of conserved structural anchors for sponge mt SSU rRNA and use them to compare 12 homologous hypervariable substructures. We define the modeled forms within each substructure as structural types and their ordered combinations across all 12 substructures as composite types. Our aims are to define the conserved scaffold, characterize the distribution of structural types and composite types among sampled lineages, and determine how compact and expanded forms are organized within a unified structure-guided coding framework.

## 2. Materials and Methods

### 2.1 Data retrieval and record filtering

Given the structural diversity of sponge mitochondrial genomes, we adopted a tiered retrieval strategy. For non-calcareous sponges, we retrieved 291 mitochondrial-genome records from NCBI, representing 232 sponge samples and including 59 RefSeq reference records. For Calcarea, we separately retrieved 15 sequences from two *Clathrina clathrus* samples and consolidated chromosome fragments corresponding to the analyzable mt SSU rRNA. In addition, four Tetractinellida samples were obtained from the Lavrov laboratory’s published resource (https://lavrovlab.github.io/Demosponge-phylogeny/published.html). After deduplication, initial sequence cleaning using NCBI2GO, and taxonomic correction against the World Porifera Database classification framework, the resulting working inventory comprised 237 candidate records. The subsequent filtering steps are summarized in Table 1 and Supplementary Table S1.

**Table 1.** Sample flow and stage-specific exclusion reasons.

| Stage | Records retained | Excluded | Stage-specific reason / note |
| --- | --- | --- | --- |
| Combined candidate inventory | 237 | — | 232 deduplicated NCBI records, one consolidated Calcarea record, and four external Tetractinellida records |
| Passed initial quality control | 236 | 1 | Non-Porifera record |
| mt SSU rRNA successfully extracted | 233 | 3 | mt SSU rRNA not detected |
| Secondary-structure models obtained | 229 | 4 | Long gaps or multiple ambiguous bases; 213 template-guided models plus 16 manually folded Hexactinellida |
| Structurally complete models | 226 | 3 | Incomplete 5' or 3' termini |
| Reliably scorable models | 219 | 7 | High structural heterogeneity prevented consistent coding |
| Final 12-substructure matrix | 216 | 3 | Class or order unresolved |

The records were filtered sequentially (Table 1 and Supplementary Table S1). One non-Porifera record was removed during initial quality control. The mt SSU rRNA was successfully extracted from 233 of the remaining 236 records; three records lacked a detectable target. Four sequences with long gaps or multiple ambiguous bases were removed before modeling, leaving 229. Models were obtained for all 229 records, 226 were structurally complete, and seven structurally heterogeneous models could not be coded consistently, leaving 219. Finally, three records lacking resolved class or order assignments were excluded. The matrix and all downstream results therefore contain 216 records only.

### 2.2 Secondary-structure modeling

Secondary-structure modeling broadly followed the comparative workflow described by Li et al. (2025). AY320032 served as the initial reference (Lavrov et al., 2005). Modeling was iterative: when an initial template produced incomplete or structurally implausible results, the record was remodeled using a more closely related lineage-specific reference selected after comparison of preliminary folds. The reference template used in the final modeling step is reported for each record in Supplementary Table S2b. The resulting structures were decomposed using bpRNA (Danaee et al., 2018) and inspected with forna visualizations (Kerpedjiev et al., 2015).

Sixteen Hexactinellida records could not be modeled reliably from the demosponge references because of low sequence homology. These records were folded manually in two stages. First, nine conserved anchors were identified and fixed to establish homologous boundaries: H9, H367, H500, H769, H885, H921, H944, H1506, and H673-loop. The first eight are stems, whereas H673-loop is a conserved loop that provides an additional anchoring point. The intervening variable regions were then folded by comparative analysis within the phylogenetic framework of Hexactinellida, considering sequence agreement, compatible base pairing, and the positions of adjacent anchors.

For each corresponding sequence group, LocARNA was used to assess whether variable regions could support reproducible secondary structures through simultaneous sequence-structure alignment and consensus folding (Will, 2024). LocARNA-supported elements were compared with the individually predicted structures. Failure to recover a stable consensus fold was recorded as an unresolved result rather than as evidence that the region lacked secondary structure. Consensus structures associated with candidate insertion regions in the 16 freshwater Spongillida records are summarized in Supplementary Table S5. All models were inspected for terminal completeness, recognizable anchors, and consistent decomposition.

### 2.3 Definition and coding of types

Twelve homologous hypervariable substructures were extracted in a fixed order: H61, H122, H240, H339, H406+H441, H577-H655, H821, H996, H1068, H1113+H1118, H1241, and H1399. The names follow the comparative rRNA convention based on homologous *E. coli* positions. The combined labels H406+H441 and H1113+H1118 are used consistently because the coded variable region spans the named component elements.

Within each specified substructure, a distinct modeled form was defined as a structural type, hereafter type. Labels A-E and, where needed, E1-E2 distinguish types within that substructure only. Thus, H61 type A and H122 type A do not imply structural equivalence. The 12 types assigned to one complete record were concatenated in the fixed order above to define its composite type. The resulting type-code string is a compact identifier for the full 12-substructure combination, not a primary-sequence alignment and not a set of nucleotide sites.

A substructure was considered variable within a taxonomic group when more than one type occurred in that group. Composite-type diversity was summarized by the number of observed composite types and Shannon entropy, H = -Σ p_i_ ln p_i_, where p_i_ is the within-group frequency of composite type i. Shannon values are descriptive and are not corrected for unequal sample sizes or sampling coverage.

### 2.4 Summary statistics

Sequence-length summaries use nucleotide counts from the structurally complete records retained in the final matrix. Counts and percentages for types use n = 216. Taxonomic summaries are reported at class, subclass, and order levels. No ancestral-state reconstruction or formal phylogenetic-signal test was performed; recurrent types are therefore described as shared or recurring rather than as independently evolved. The taxonomic relationships displayed in Figs. 1 and 2 were redrawn from the topology summarized and, for Demospongiae, inferred by Lavrov et al. (2023). Only the branching order was retained; branch lengths, node spacing, and terminal positions were adjusted for display and have no quantitative meaning. These diagrams were used to organize the sampled taxa visually and were not inferred from the mt SSU rRNA data analyzed here.

**Figure 1.**
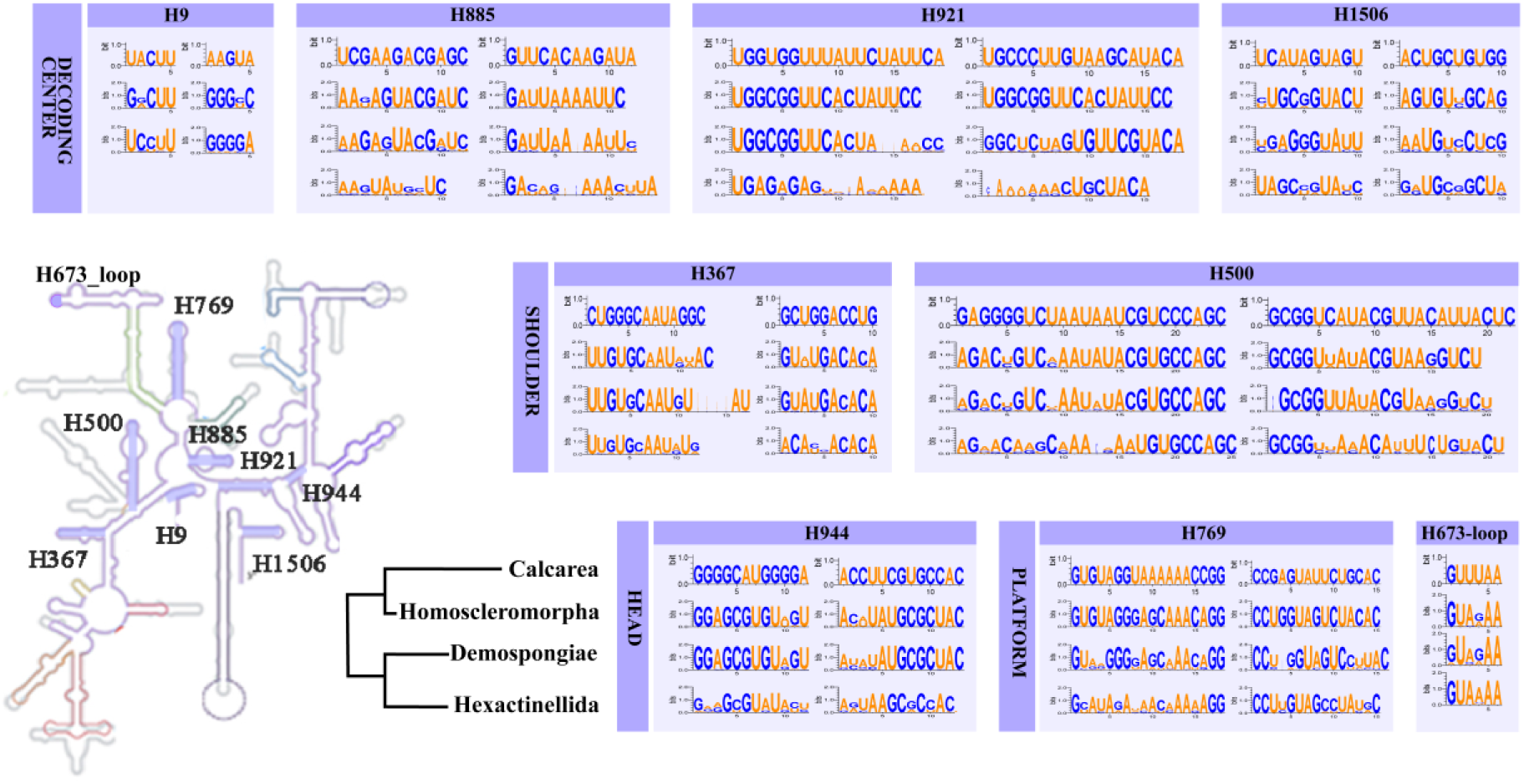
Nine conserved structural anchors in sponge mt SSU rRNA. Eight anchors are stems (H9, H367, H500, H769, H885, H921, H944, and H1506); H673-loop is a conserved loop and provides an additional anchoring point. Sequence logos summarize representatives of the four sampled classes. The class-level topology follows the relationships summarized by Lavrov et al. (2023) and is shown schematically; branch lengths are not to scale.

**Figure 2.**
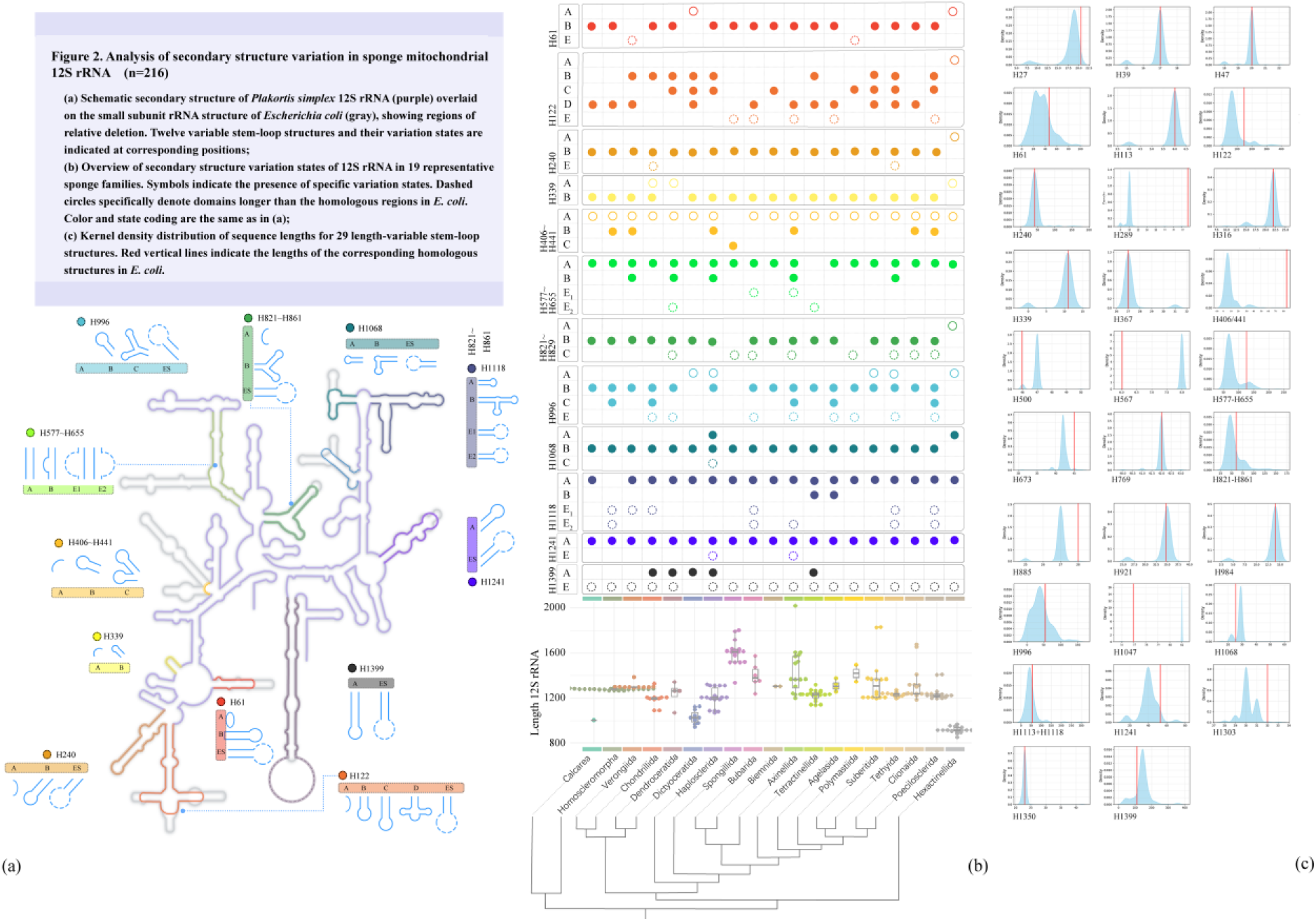
Structural variation in sponge mitochondrial SSU rRNA. (a) Secondary-structure map showing the 12 hypervariable substructures and their structural types. (b) Distribution of types and mt SSU rRNA lengths across sampled sponge groups (n = 216); dashed circles indicate candidate insertion regions. (c) Length distributions of 29 variable structural elements; red lines indicate the corresponding *E. coli* homolog lengths. Taxonomic groups are arranged according to the relationships summarized and, for Demospongiae, inferred by Lavrov et al. (2023). Branch lengths are schematic and are not to scale.

## 3. Results

### 3.1 Final dataset and length variation

The final matrix contained 216 records from four classes and 22 named orders (Supplementary Tables S2a-S2c and Supplementary Data S6). Demospongiae contributed 177 records, Homoscleromorpha 22, Hexactinellida 16, and Calcarea one. The largest orders were Homosclerophorida (n = 22), Tetractinellida (n = 19), Haplosclerida (n = 18), and Axinellida, Spongillida, and Suberitida (n = 16 each). All analytical tables and percentages exclude the three records with unresolved class or order assignments.

The mt SSU rRNA length ranged from 853 to 2,019 nt (mean 1,269.8 nt; SD 190.8). Demospongiae ranged from 941 to 2,019 nt, Hexactinellida from 853 to 969 nt, and Homoscleromorpha from 1,267 to 1,297 nt; the single Calcarea record was 1,003 nt. The sampled Spongillida records were long (1,516-1,801 nt; mean 1,618.9 nt; SD 88.4), whereas the hexactinellid records were consistently shorter (mean 916.6 nt; SD 28.5).

### 3.2 Conserved structural anchors

Nine conserved structural anchors were identifiable across the four sampled classes (Fig. 1). Eight anchors are stems: H9, H367, H500, H769, H885, H921, H944, and H1506. H673-loop is the ninth anchor. Although it is a loop rather than a stem, its positional and sequence conservation made it useful for fixing local homology. The anchors occupy the decoding-center, head, platform, and shoulder regions of the modeled small subunit.

These anchors were used as methodological reference points, not as an experimentally demonstrated minimal functional core. Their consistent recovery enabled cross-class comparison and was essential for the manual reconstruction of Hexactinellida. Calcarea is represented by one record, so apparent cross-class conservation should be reevaluated as additional calcareous sponge data become available.

### 3.3 Types across the 12 hypervariable substructures

Across the 12 hypervariable substructures, 38 types were observed (Fig. 2a; Supplementary Tables S2c and S3). Diversity was strongly uneven. H122 contained five types; H577-H655, H996, and H1113+H1118 contained four each. H61, H240, H406+H441, H821, and H1068 contained three each, whereas H339, H1241, and H1399 contained two each.

Several substructures were dominated by one type: H1241 type A occurred in 210 records (97.2%), H1068 type B in 196 (90.7%), H240 type B in 189 (87.5%), H406+H441 type A in 188 (87.0%), H339 type B in 186 (86.1%), and H1399 type E in 185 (85.6%). H122 was the most even, with its most common type B occurring in 65 records (30.1%); H996 type B occurred in 110 (50.9%). These frequency differences identify substructures with broadly shared forms and others that contribute disproportionately to observed structural diversity.

### 3.4 Composite types and order-level diversity

The ordered combinations across the 12 substructures yielded 62 composite types (Fig. 2b; Supplementary Tables S2c and S4). Demospongiae contained 58 composite types among 177 records and was variable at all 12 substructures. Heteroscleromorpha contained 43 composite types among 142 records, Keratosa nine among 15, and Verongimorpha eight among 20. Homoscleromorpha contained three composite types among 22 records. Hexactinellida and Calcarea each had one observed composite type, although Calcarea was represented by only one record.

At the order level, Haplosclerida and Clionaida each contained nine composite types; Axinellida and Suberitida each contained seven. Their Shannon indices were 2.043, 1.899, 1.787, and 1.661, respectively. These values describe the observed frequency distributions but are not standardized comparisons of evolutionary rate. The most frequent composite types in the complete dataset were BDBBAABBBAAE (n = 21), BCBBAABBBAAE (n = 20), BEBBAE1EEBAAE (n = 17), and AAAAAAAAAAAE (n = 16).

### 3.5 Contrasting lineage-associated patterns

All 16 Hexactinellida records shared composite type AAAAAAAAAAAE, which combines type A at the first 11 substructures and type E at H1399. All 16 sampled Spongillida records shared composite type BEBBAE1EEBAAE. The two lineages were therefore both uniform in the coded matrix despite markedly different sequence lengths. Structural uniformity within a sampled lineage is not equivalent to structural compactness.

Structure-guided comparison identified four to five candidate insertion regions in each of the 16 sampled freshwater Spongillida records (Supplementary Table S5), whereas corresponding candidate insertion regions were not detected in the analyzed marine records. Comparable H7/H8-region insertions in Lake Baikal sponges have been reported to form hairpin-like elements (Lavrov, 2010; Lavrov et al., 2012). Some candidate regions in the current models did not contain predicted hairpins under the applied criteria; this weakens their classification as hairpin-containing regions but does not by itself negate the underlying inserted sequence. Because freshwater habitat and lineage are confounded, the observed distribution cannot distinguish environmental adaptation from inheritance within the sampled freshwater lineage.

## 4. Discussion

### 4.1 A conserved scaffold accommodates localized variation

The nine conserved anchors provided stable reference points for comparing 216 taxonomically resolved records, including highly divergent Hexactinellida, within a common structure-guided framework. These anchors are best interpreted as a conserved structural scaffold rather than as a demonstrated minimal functional core. Their consistent recovery across the sampled records supports the persistence of a shared scaffold, while the 12 hypervariable substructures locate the principal differences among the modeled structures.

Variation was unevenly distributed across substructures and lineages. Several substructures were strongly dominated by one type, whereas H122, H577-H655, H996, and H1113+H1118 showed greater type diversity and are therefore priorities for evaluating covariation support and alternative folds. Demospongiae, particularly Heteroscleromorpha, accounted for most composite types, but this pattern is inseparable from uneven sampling because larger groups provide more opportunities to recover rare combinations. Group-level richness and Shannon indices should therefore be interpreted as descriptions of the present dataset rather than as estimates of intrinsic evolutionary rates. The matrix supports the recognition of lineage-associated structural profiles, but distinguishing inheritance from convergence, reversal, or similarity introduced by categorical coding will require explicit character mapping and tests of phylogenetic signal.

### 4.2 Overall size and composite type describe distinct dimensions of structural variation

Overall length and the composition of the 12 coded substructures capture different aspects of mt SSU rRNA structure. This distinction is illustrated by the sampled Hexactinellida and freshwater Spongillida: each lineage was uniform for a different composite type, yet they differed markedly in total length and local structure. Because a composite type summarizes only the predefined substructures, it should not itself be classified as compact or expanded; those descriptors apply to the complete modeled molecule.

The coexistence of compact and highly expanded extant structures broadens the empirical range that constructive-reductive models of mitoribosome evolution must explain (van der Sluis et al., 2015). It does not, however, identify the ancestral condition or the direction of structural change. Such inferences will require mapping structural characters onto a well-supported species tree.

With the exception of Hexactinellida, all sampled sponge classes contained at least some local regions that exceeded the length of their homologues in *E. coli*. This pattern was especially pronounced in freshwater Spongillida, in which the candidate insertion sequences also showed clear homology across records. Nevertheless, sequence alignment and related analyses revealed no features supporting their annotation as introns. We therefore treat these regions as lineage-associated components of the modeled mt SSU rRNA rather than as intronic elements. Although their consistent occurrence in freshwater Spongillida is notable, the present evidence is insufficient to interpret these structural features as adaptations to freshwater environments.

### 4.3 Methodological scope, limitations, and prospective applications

The conserved-anchor workflow and type matrix make positional-homology decisions explicit and auditable. The coding system deliberately reduces complex modeled structures to categorical characters: a type denotes a distinct modeled form only within its specified substructure, and a composite type cannot substitute for inspection of the complete fold. Template-guided prediction, manual comparative curation, categorical coding, and uneven taxonomic sampling remain important sources of uncertainty. The matrix therefore supports comparative description and hypothesis generation but does not by itself establish function, ancestral states, or adaptation.

The framework suggests two practical directions for further work. First, by delimiting comparable target regions, it may inform evaluation of mt SSU rRNA as a complementary molecular marker. However, structural conservation of the anchors does not ensure primary-sequence conservation or primer performance, and amplification success, eDNA recovery, and taxonomic resolution were not tested here (Rossouw et al., 2024; Timmers et al., 2022). Second, the type matrix provides discrete characters for ancestral-state reconstruction and phylogenetic-signal analysis, while the underlying alignments and folds permit covariation tests, matched nuclear-mitochondrial comparisons, and experimental evaluation of candidate insertions. These applications can be pursued without treating the observed differences as evidence of directional change, independent origins, or functional necessity.

## 5. Conclusion

Using nine conserved structural anchors, this study placed 216 taxonomically resolved sponge mt SSU rRNA records from four classes and 22 orders into a common comparative framework. The 12 hypervariable substructures contained 38 structural types and formed 62 composite types, while sequence length ranged from 853 to 2,019 nt. The central contribution is the structure-guided coding framework: it converts rRNAs that are difficult to compare by primary sequence into homologous, auditable structural characters and separates broadly conserved elements from localized variation.

The matrix documents the coexistence of compact and extensively expanded structures among sponge lineages. Hexactinellida and sampled freshwater Spongillida were each uniform for a different composite type despite their sharply contrasting overall structures. These observations support lineage-associated structural organization within a shared scaffold but do not establish ancestral states, directionality, independent origins, adaptation, or phylogenetic signal. The models, definitions, and record-level matrix provide a reusable resource for testing those hypotheses and for evaluating future structure-guided markers.

## Acknowledgements

We thank Dennis V. Lavrov for valuable comments on the study and for discussions that helped refine our interpretation of sponge mitochondrial rRNA evolution. The authors thank the members of the Lab of Insect Systematics and Evolutionary Biology (LISEB), Jiangxi Normal University, for their contributions to data curation, database development, and technical support. This work was supported by the National Natural Science Foundation of China (Grant Nos. 32370500, 42576137 and 62662044), and the Shandong Provincial Natural Science Foundation (Grant No. ZR2023MD100).

## Data Availability

All data underlying this study are provided in the accompanying supplementary materials. Supplementary Table S1 contains the 237-record candidate inventory and stage-specific exclusion decisions. Supplementary Table S2a summarizes the taxonomic composition of the final 216-record dataset; Supplementary Table S2b provides record-level sequences, structural annotations, segment measurements, and final reference-template accessions; and Supplementary Table S2c provides structural-type assignments for the 12 hypervariable substructures and the corresponding composite types. Supplementary Table S3 summarizes structural-type diversity by substructure, Supplementary Table S4 reports order-level composite-type diversity, and Supplementary Table S5 provides consensus features of candidate insertion regions in sampled freshwater Spongillida. Supplementary Data S6 contains the 216 predicted secondary structures in STR format.

